# SpliceCoord: A computational toolkit to identify coordinated exon splicing events from long-read transcriptome sequencing data

**DOI:** 10.1101/2025.08.20.671187

**Authors:** Pallavi Gupta, Ishaan Gupta

## Abstract

As long-read RNA sequencing technologies emerge and improve, they offer a greater depth and precision of the transcriptome of organisms, tissues, and cells, enabling deeper exploration of alternative splicing patterns. It has been observed that specific pairs of exons appear together in a higher or lower frequency than is expected by chance. Such a mutual inclusion or exclusion may reflect underlying structural or functional constraints; however, a systemic method for this analysis is lacking. Here, we develop an R-based computational toolkit, called *SpliceCoord*, to identify coordinated splicing of exons using a transcript-wise count matrix and a GTF file that maps the expressed transcripts and their exons to the genome.

## INTRODUCTION

In eukaryotes, splicing is the process by which the fragments of a gene called introns are removed from the pre-mRNA and the remaining segments, known as exons, are stitched together, forming a mature mRNA. Alternative splicing allows exons to be assembled in different combinations, without changing the order, leading to a much larger number of RNA and protein isoforms than that of genes [1, 2]. However, this assembly of exons is non-random, i.e., certain exons may be preferentially spliced in or out together, through a phenomenon called coordinated splicing. Two forms of coordinate splicing can be observed – mutual inclusion and exclusion of exons. In the former, two exons appear together in transcripts of a gene more often than can occur if it were a random event. In contrast, the frequency of occurrence of both exons in the same transcript is lower than expected for the latter. While tools such as *rMATS, SUPPA2, FLAIR*, and *SQANTI3* study mutual exclusion of exons from short and long-read sequencing data, it is not complemented by detecting co-inclusion events [3–6].

A handful of studies have explored the idea of coordinated splicing [7–9]; however, no systematic detection pipeline exists. Here, we describe a simple and intuitive R-based toolkit that uses transcript counts and a GTF file that maps the relevant genes, transcripts, and their exons to the genome as inputs and returns a tab-separated file with metrics that quantify the extent of coordination of pairs of exons.

## METHODS

To demonstrate the workflow, we use a small, manually curated dataset consisting of 10 genes and 25 transcripts (Figure 1 and Supplementary Table 1). All exons are colour-coded with respect to their genes, and the introns are drawn with thin lines connecting them. While the exons have been drawn to indicate their relative lengths, this is not true for introns here. The numbers next to gene and transcript identity are the full-length counts of that isoform. A genome GTF file corresponding to this dataset with entries for genes, transcripts, and exons was also created (Table 1). Here, the genes have been given a varying number of exons, ensuring enough combinations of constitutive and alternative exons are generated. For simplicity, each gene is placed in its own chromosome, and all exon lengths are multiples of three, as here we are not interested in whether the resulting protein from these isoforms will be coding or not.

**Table 1.**
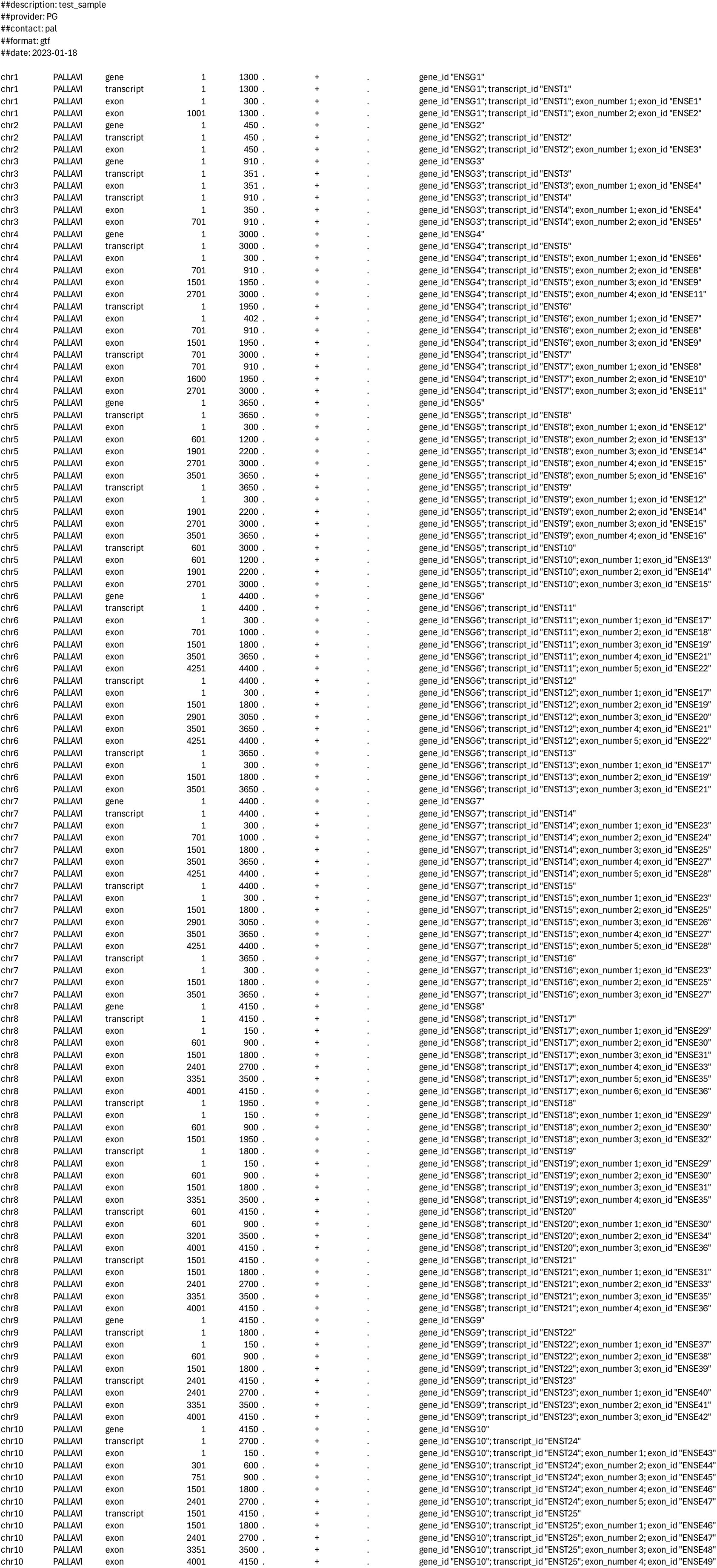
Reference GTF file created for the curated dataset.

**Figure 1.**
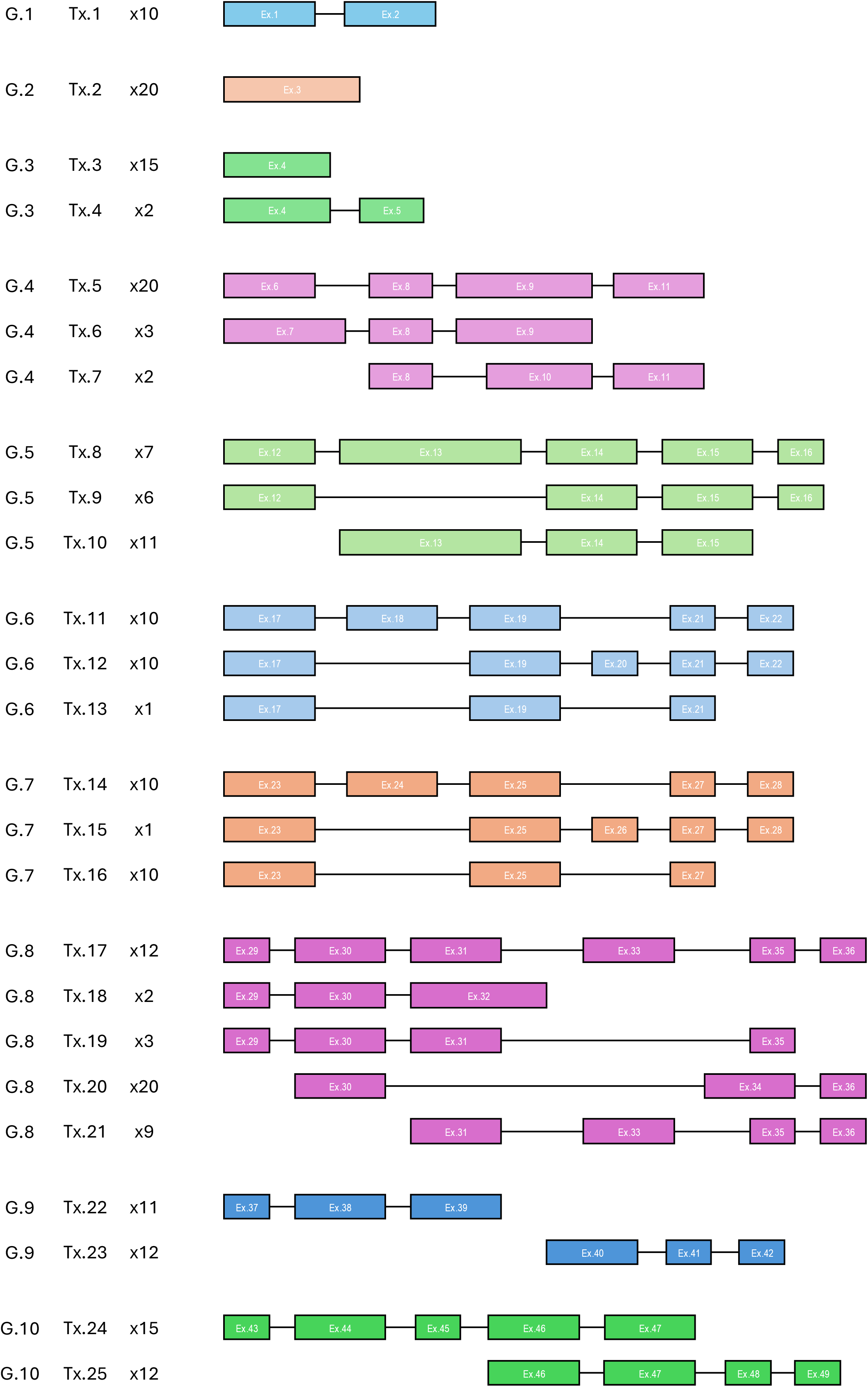
Curated dataset with 10 genes (G.1 – G.10), 25 transcripts (T.1 – T.25), and 49 exons (Ex.1 – Ex.49) with the transcript counts. Each color corresponds to one gene.

Inputs from the user

1. Counts of long-read outputs with two columns named ‘*isoform’* and ‘*counts*’

2. Genome GTF corresponding to transcripts in the counts file must have genomic coordinates for genes, transcripts, and exons. If the user has a transcriptome GTF obtained after reconstruction from *FLAIR* [5] or *Stringtie2* [10], additional tools must be used to modify that to a script-compatible format.

3. One tab-separated index file named ‘sample_names’ with three columns but no column headers. The first column is a short-hand notation of the samples (to be used as a prefix for outputs of all steps), the second column is the filename of the count matrix (including the extension, if any), and the third column is the filename of the genome GTF file (including the extension, if any).

Pipeline –

Step 0 – All the requisite R packages are checked and, if not found, installed. These include CRAN packages *dplyr, tibble, parallel, BiocManager*, as well as Bioconductor packages *rtracklayer* and *GenomicFeatures*. We recommend creating a conda environment with a recent R version to run this script.

Step 1 – First, the user-supplied genome GTF file is converted to a *TxDb* object, which is used to create exon- and transcript-wise index files, including renaming exons to the format <GENEID>_<start>_<end> to deal with redundant exons that have different IDs. This is the most time-consuming step in the entire pipeline, where multiple cores can benefit the runtime.

Step 2 – This has some of the most crucial filtration steps that eliminate lowly expressed genes, truncated transcripts, and spurious exons that can alter the results if not handled correctly.

- We begin by filtering out genes with a single transcript – <u>this removes genes G.1 and G.2</u>
- In a typical long-read workflow, the full-length counts may not be too high. Therefore, it is essential to use a cutoff. We have chosen an arbitrary cutoff of 10 reads from all transcripts of the gene combined.
- Next, any spurious exons, which are supported by less than 5% of the total reads from that gene, are eliminated – <u>this removes exons Ex.26 and Ex.32</u>
- All genes must contain three or more – <u>this removes gene G.3</u>
- As can be observed for gene G.4, exons 1 and 2, and exons 4 and 5 overlap, but have at least one of the splice sites different. Generally, this difference in length is much shorter than the distance between adjacent exons; hence, we have collapsed any such exons sharing the 5’ or 3’ splice site. These are referred to as ‘exon chunks’
- Again, any gene with up to two exon chunks is removed from further analysis.
- In some cases, when a large number of 5’ and 3’ truncations are obtained, most or all such fragments are non-overlapping. Unless they share at least one exon, these are discarded as they do not represent real molecules – <u>this removes transcripts Tx.22, Tx.23 from gene G.9</u>
- Another possibility is where the start of one is the same as the end of another, or vice-versa, these must be omitted as well – <u>this removes transcripts Tx.6, Tx.7 and Tx.24, Tx.25 from genes G.4 and G.10, respectively</u>
- One final check is done to exclude any remaining genes with just one transcript – <u>this removes Tx.5 of G.4</u>

Step 3 – Next, we identify transcripts that may be fragments of a longer full-length (FL) transcript. This is done via determining the transcription start site (TSS or the first exon chunk of that gene) and the transcription termination site (TTS or the last exon chunk of that gene). These truncations (T) are not removed, just marked as ‘FL’ or ‘T’ for calculations, see Tx.10 and Tx.21 in Figure 2. If only one true FL transcript is identified for a gene, then the gene is eliminated.

**Figure 2.**
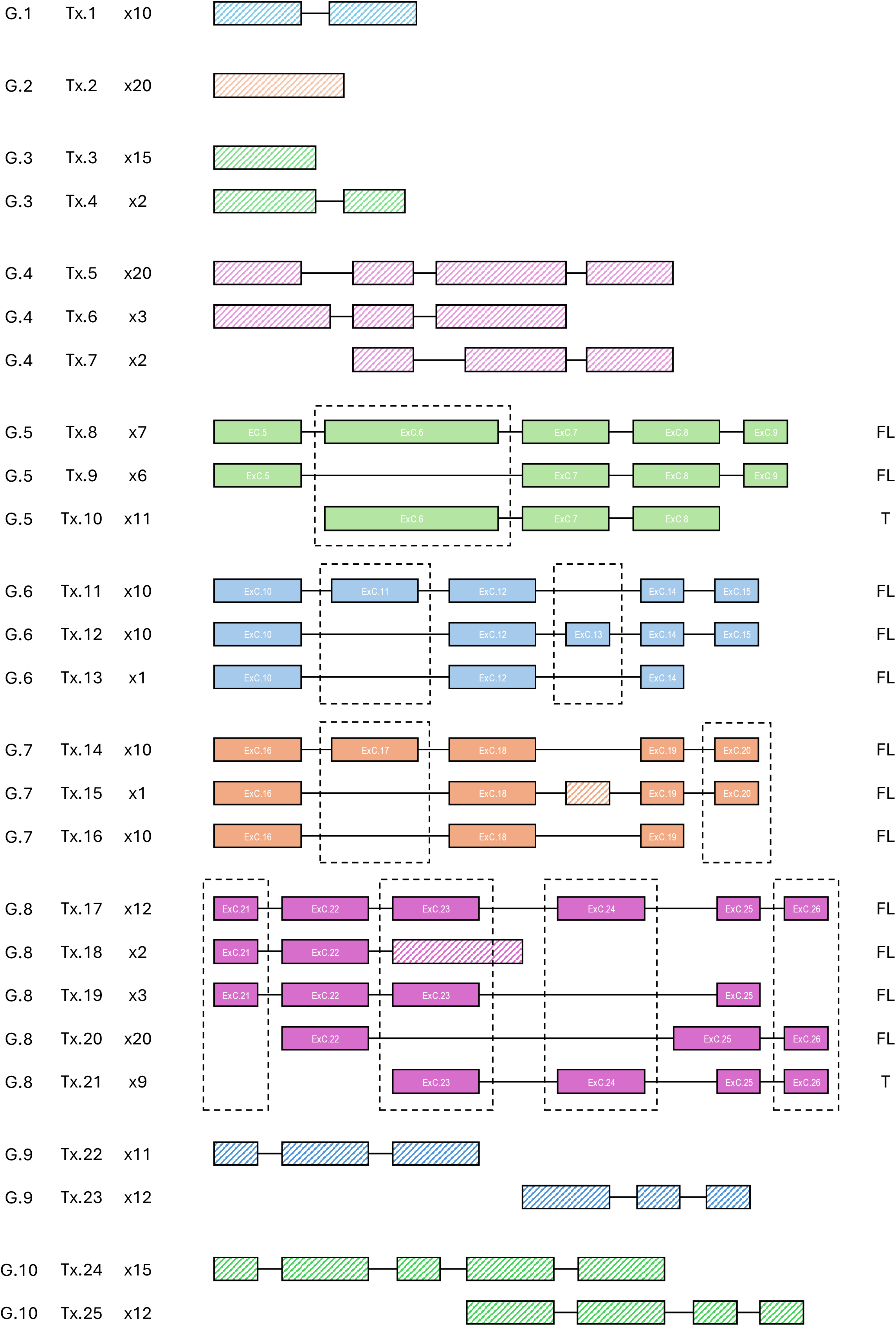
The final 22 exon chunks (ExC.5 – ExC.26) detected in the curated dataset after the cutoffs and filters. FL vs T marks the transcripts’ full-length vs truncated status. The dashed lines show all alternative exon chunks.

Step 4 – Constitutive and alternative exon chunks are identified as those present in >95% and <95% reads that contain those exon chunks, respectively. Here, the demarcation of T vs FL transcripts is used – while all the FL transcripts are considered for all exon chunks, those marked as T are only used if the exon chunks that fall within the range of TSS and TTS of that transcript. All alternative exon chunks are outlined by broken lines in Figure 2.

Step 5 – Pairs are made for all the identified alternative exon chunks per gene – any gene with only one alternative exon chunk will not be a part of the downstream steps – <u>such as ExC.6 of gene G.5</u>. Each exon chunk is labelled as ‘end or ‘internal’ based on whether it is at the terminals or not. They are also categorized as ‘adjacent’ or ‘distant1’ or ‘distant2’ exon chunks if they are next to each other or have at least one separating exon chunk, or have at least one constitutive exon between them, respectively.

Step 6 – This is where calculations are done to determine the pattern, if any, and the direction of coordinates splicing of a pair of exon chunks 1 and 2.

- FL transcripts that contain each of the exon chunks, and T transcripts that must be considered for each (with the same logic as in step 4), are identified. Next, four values are calculated: *a* is the number of molecules with both exon chunks,*b* is the number of molecules with exon chunk 1 but not 2, *c* is the number of molecules with exon chunk 2 but not 1, and is the number of molecules that contain neither 1 nor 2.
- Three values, σ (sigma), σ’(sigma-prime), and (co-inclusion score), are calculated as follows–

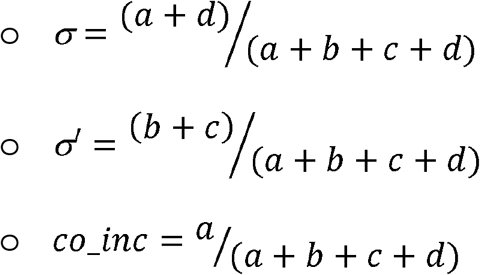
- Fisher’s test is performed, and the p-value is noted. Multiple testing correction is performed per category, adjacent, distant1, and distant2, using the Benjamini– Yekutieli (BY) method [11]
- Finally, the log odds ratio (LOR) is calculated as 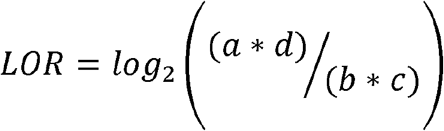; if any value is zero, it is instead changed to 0.5

Once the tabulation is complete, significantly coordinated pairs of exon chunks are identified based on corrected p-value < 0.05. Mutually inclusive pairs have, *LOR > 0, σ > 0.7, σ’< 0.3* and co _inc_ > 0.3; while mutually exclusive pairs have. *LOR* < 0, σ < 0.3, and σ’ > 0.7. The pipeline returns a tab-separated file per sample as the final output. In addition, and all the intermediate steps have associated RDS objects that can be used to follow through the filtration steps.

All example files to reproduce the results are provided in the GitHub documentation. We recommend running *SpliceCoord* on a server/HPC because the intermediate files can be large. The seven steps, including installation, can be run individually or in one go using *wrapper_script.sh*.

Individual commands –

*Rscript step0_install.R*

*Rscript step1_txdb_obj.R arg1*

output → txdb_<sample>; exon_<sample>.RDS; tx_<sample>.RDS

*Rscript step2_exonchunks.R arg2*

output → <sample>_used.RDS; updated → exon_<sample>.RDS; tx_<sample>.RDS

*Rscript step3_TSS_TTS.R*

output → exon_<sample>_final.RDS;tx_<sample>_final.RDS

*Rscript step4_const_alt_exons.R*

output → constitutive_EXONCHUNK_<sample>.RDS;

non_constitutive_EXONCHUNK_<sample>.RDS

*Rscript step5_pairs.R arg2*

output → Table_<sample>.RDS

*Rscript step6_calculations.R arg2 arg3*

output → filled_<sample>.tsv; filled_<sample>.RDS; significant_<sample>.RDS

Single command version –

*bash wrapper_script.sh arg1 arg2 arg3*

where arg1 is the number of threads while generating two index files for each genome GTF used in step1 (default takes one-fourth of the available threads); arg2 is the number of threads for in subsequent steps 2, 5, and 6 (default takes one-eighth of the available threads); and arg3 can be ‘*annot*’ if the genome GTF file contains a ‘*gene_name*’ column and the user wishes to add that information to the final output .csv file, or arg3 can be any other word.

Testing on the SG-NEx dataset

Minimap2 [12], FLAIR [5], and SQANTI3 [6] were used to obtain the counts and GTF files corresponding to the transcriptome, which was then converted to a genome-GTF format using *gffread [13], agat* (https://github.com/NBISweden/AGAT), and *sed*. An index file was manually written, and the run_wrapper.sh script was used to test the pipeline.

## RESULTS

Five out of eight tested exon chunk pairs from three genes show coordinated splicing for the manually curated dataset. These include mutually exclusive pairs ExC.11 and ExC.13 from gene G.6 and mutually inclusive pairs ExC.17 and ExC.20 from G.7. Additionally, from G.8, exon chunks ExC.21, ExC.23, and ExC.24 all show mutual inclusion with each other. At the same time, none of them have significantly coordinated splicing with the remaining exon chunk ExC.26. All the tested pairs and their coordination metrics are provided in Table 2.

**Table 2.**
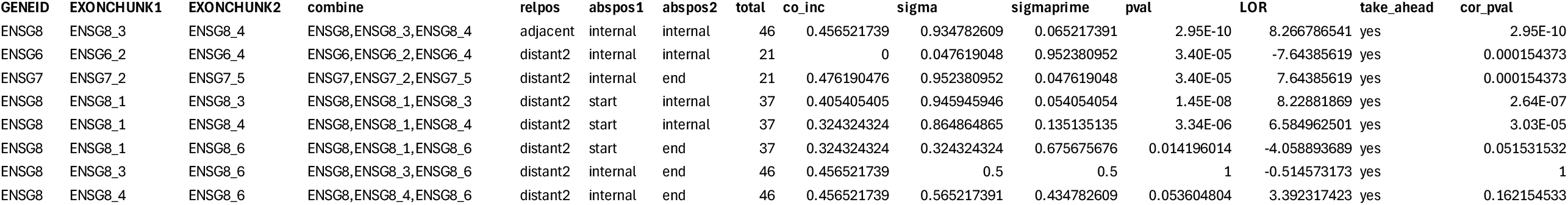
*SpliceCoord* output for the curated dataset.

We additionally generated five mock datasets (Supplementary Tables 2-6) that preserved the approximate ratios of transcript abundances as used earlier but varied in total counts (200k, 100k, 50k, 10k, and 1k). Across them all, the same five exon chunk pairs were consistently identified as significantly coordinated, demonstrating the robustness of our pipeline.

We also tested the pipeline on Nanopore direct cDNA-seq data from six cell lines, including five cancer lines, A549, Hct116, HepG2, K562, and MCF7, and one embryonic stem cell line, H9 [14]. After long-read alignment, quantification, and classification, the counts were used directly, while the GTF file was transformed to the script-compatible format, as outlined in *Methods*. Table 3 summarizes of the number of exon chunks tested and their coordination. While the absolute number of exon pairs that can be tested for coordination generally increases with a higher sequencing depth, due to more unique isoforms and their counts, this is not always true, as evident for Hct116 (Figure 3A). Similarly, owing to innate trends in isoform arrangement, the coordinated pairs may fall into the three categories, ‘adjacent’, ‘distant1’, or ‘distant2’, in different ratios (Figure 3B).

**Table 3.**
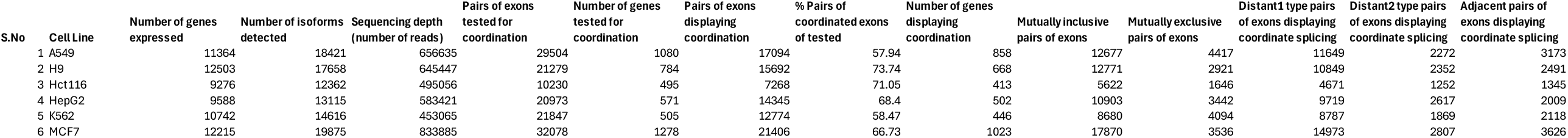
Summary of the *SpliceCoord* output for the SG-NEx dataset.

**Figure 3.**
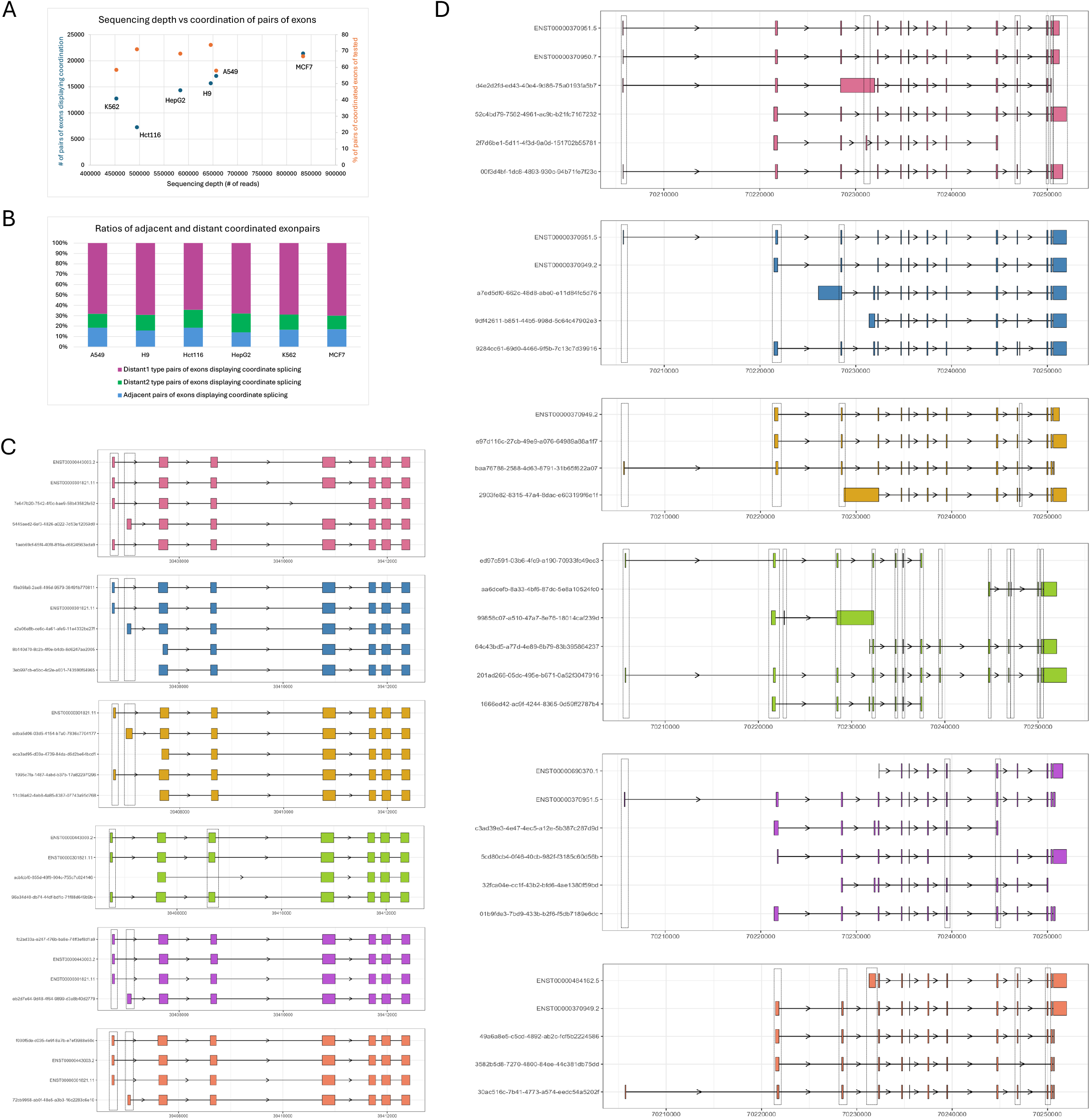
Summary of the SG-NEx cDNA long-read sequencing dataset’s coordinated splicing. A, Sequencing depth vs number of pairs of exons displaying coordination (left axis) and vs percentage of pairs of coordinated exons of the total pairs tested (right axis). B, Percentage of adjacent, distant1, and distant2 type of coordinated exon pairs per cell line. Isoforms for RPSA (C), and SRSF11 (D) that passed the filters and were tested for coordination in cell lines in the order A549, H9, Hct116, HepG2, K562, and MCF7. Dashed lines indicate alternative exon chunks that show at least one of either type of coordination with another such exon chunk.

We demonstrate the coordination in splicing for two genes – *RPSA* (ribosomal protein SA, ENSG00000168028) and *SRSF11* (serine arginine rich splicing factor 11, ENSG00000116754). RPSA has four to five isoforms made of seven or eight exon chunks, while *SRSF11* had four or six isoforms comprising of a total of fourteen to sixteen exon chunks (Supplementary Table 7). The splicing coordination patterns for *RSPA* are consistent across all five of the six cell lines, with one adjacent exon chunk pair displaying mutually exclusive coordinated splicing, while for HepG2, the exon chunks 1 and 3 display mutual inclusion (Figure 3C). On the other hand, a considerable variation in terms of coordinated pairs of exon chunks can be observed for *SRSF11*, owing to a variation in the number of alternatively spliced exons across the cell lines (Figure 3D).

## DISCUSSION

The concept of coordination of expression and splicing of multiple genes in concert with cell states or tissue identity is prevalent [15, 16]. However, the coordination of the splicing of adjacent or distant exons of a gene is not appreciated. While the exact cause and mechanism of coordinated splicing of exons is unknown, their mutual inclusion or exclusion can arise from structural or functional constraints.

An increased use of long-read RNA and cDNA sequencing technologies, both in bulk and single-cell contexts [17, 18], has the potential to uncover diverse splicing patterns. *SpliceCoord* is the first publicly available method designed to robustly detect coordinated splicing events from such transcript count data, thereby enabling better quantification of their abundance and prevalence.

## DATA AVAILABILITY

No new data was generated in this study

## CODE AVAILABILITY

All the codes and example files used in this study are available on GitHub (https://github.com/pgupta3005/SpliceCoord)

## FIGURE AND TABLE LEGENDS

Supplementary Table 1: Curated dataset

Supplementary Table 2: Curated dataset with 200,000 reads

Supplementary Table 3: Curated dataset with 100,000 reads

Supplementary Table 4: Curated dataset with 50,000 reads

Supplementary Table 5: Curated dataset with 10,000 reads

Supplementary Table 6: Curated dataset with 1000 reads

Supplementary Table 7: Summary of RPSA and SRSF11 genes from *SpliceCoord*

## REFERENCES

1. Wright, C.J., C.W.J. Smith, and C.D. Jiggins, Alternative splicing as a source of phenotypic diversity. Nat Rev Genet, 2022. 23(11): p. 697–710.

2. Marasco, L.E. and A.R. Kornblihtt, The physiology of alternative splicing. Nat Rev Mol Cell Biol, 2023. 24(4): p. 242–254.

3. Shen, S., et al., rMATS: robust and flexible detection of differential alternative splicing from replicate RNA-Seq data. Proc Natl Acad Sci U S A, 2014. 111(51): p. E5593–601.

4. Trincado, J.L., et al., SUPPA2: fast, accurate, and uncertainty-aware differential splicing analysis across multiple conditions. Genome Biol, 2018. 19(1): p. 40.

5. Tang, A.D., et al., Full-length transcript characterization of SF3B1 mutation in chronic lymphocytic leukemia reveals downregulation of retained introns. Nat Commun, 2020. 11(1): p. 1438.

6. Pardo-Palacios, F.J., et al., SQANTI3: curation of long-read transcriptomes for accurate identification of known and novel isoforms. Nat Methods, 2024. 21(5): p. 793–797.

7. Tilgner, H., et al., Comprehensive transcriptome analysis using synthetic long-read sequencing reveals molecular co-association of distant splicing events. Nat Biotechnol, 2015. 33(7): p. 736–42.

8. Tilgner, H., et al., Microfluidic isoform sequencing shows widespread splicing coordination in the human transcriptome. Genome Res, 2018. 28(2): p. 231–242.

9. Hardwick, S.A., et al., Single-nuclei isoform RNA sequencing unlocks barcoded exon connectivity in frozen brain tissue. Nat Biotechnol, 2022. 40(7): p. 1082–1092.

10. Kovaka, S., et al., Transcriptome assembly from long-read RNA-seq alignments with StringTie2. Genome Biol, 2019. 20(1): p. 278.

11. Benjamini, Y. and D. Yekutieli, The control of the false discovery rate in multiple testing under dependency. The Annals of Statistics, 2001. 29(4): p. 1165-1188, 24.

12. Li, H., Minimap2: pairwise alignment for nucleotide sequences. Bioinformatics, 2018. 34(18): p. 3094–3100.

13. Pertea, G. and M. Pertea, GFF Utilities: GffRead and GffCompare. F1000Res, 2020. 9.

14. Chen, Y., et al., A systematic benchmark of Nanopore long-read RNA sequencing for transcript-level analysis in human cell lines. Nat Methods, 2025. 22(4): p. 801–812.

15. Fagnani, M., et al., Functional coordination of alternative splicing in the mammalian central nervous system. Genome Biol, 2007. 8(6): p. R108.

16. Baralle, F.E. and J. Giudice, Alternative splicing as a regulator of development and tissue identity. Nat Rev Mol Cell Biol, 2017. 18(7): p. 437–451.

17. Ament, I.H., et al., Long-read RNA sequencing: A transformative technology for exploring transcriptome complexity in human diseases. Mol Ther, 2025. 33(3): p. 883–894.

18. Gupta, P., et al., Advances in single-cell long-read sequencing technologies. NAR Genom Bioinform, 2024. 6(2): p. qae047.

